# Interaction Between Phosphoinositide 3-Kinase and the VDAC2 Channel Tethers Endosomes to Mitochondria and Promotes Endosome Maturation

**DOI:** 10.1101/2021.01.18.427063

**Authors:** Aya O. Satoh, Yoichiro Fujioka, Sayaka Kashiwagi, Aiko Yoshida, Mari Fujioka, Hitoshi Sasajima, Asuka Nanbo, Maho Amano, Yusuke Ohba

**Author notes:** Center for Advanced Research and Education, Asahikawa Medical University, 2-1-1-1 Midorigaoka Higashi, Asahikawa 078-8510, Japan (Hitoshi Sasajima). Research Center for the Control and Prevention of Infectious Diseases, Nagasaki University, 1-12-4 Sakamoto, Nagasaki 852-8523, Japan (Asuka Nanbo).

## Abstract

Intracellular organelles of mammalian cells communicate with each other during various cellular processes. The functions and molecular mechanisms of such interorganelle association remain largely unclear, however. We here identified voltage-dependent anion channel 2 (VDAC2), a mitochondrial outer membrane protein, as a binding partner of phosphoinositide 3-kinase (PI3K), a regulator of clathrin-independent endocytosis downstream of the small GTPase Ras. VDAC2 was found to tether endosomes positive for the Ras-PI3K complex to mitochondria in response to cell stimulation with epidermal growth factor and to promote clathrin-independent endocytosis as well as endosome maturation at membrane contact sites. With a newly developed optogenetics system to induce mitochondrion-endosome association, we found that, in addition to its structural role in such association, the pore function of VDAC2 is also required for the promotion of endosome maturation. Our findings thus uncover a previously unappreciated role of mitochondrion-endosome association in the regulation of endocytosis and endosome maturation.

**Highlights:**

- The mitochondrial protein VDAC2 binds PI3K and tethers endosomes to mitochondria
- VDAC2 promotes clathrin-independent endocytosis
- VDAC2-PI3K interaction induces acidification of endosomes associated with mitochondria
- The pore function of VDAC2 also contributes to endosome maturation at contact sites

## INTRODUCTION

Endocytosis mediates the internalization of plasma membrane proteins such as receptors as well as the ingestion of a variety of exogenous nutrients and pathogens in eukaryotic cells. Cargo incorporated in endocytic vesicles is transported to early endosomes and is then sorted during subsequent fusion events between endosomes. Interendosomal fusion is regulated by the small GTPase Rab and soluble *N*-ethylmaleimide–sensitive factor (NSF) attachment protein receptor (SNARE) proteins (Langemeyer et al., 2018). For example, vacuolar protein sorting 34 (Vps34), a class III phosphoinositide 3-kinase (PI3K), is recruited to and activated at early endosomes by Rab5, resulting in the generation of phosphatidylinositol 3-phosphate [PI(3)P]. Early endosome antigen 1 (EEA1) then binds to both Rab5 and PI(3)P in the endosome membrane through its FYVE domain and thereby tethers early endosomes together with SNARE proteins (Christoforidis et al., 1999). The fates of material loaded in early endosomes include degradation as a result of its transfer to lysosomes or recycling to the plasma membrane via recycling endosomes. In the process of endocytic degradation, early endosomes undergo maturation into late endosomes in association with a gradual decrease in intravesicular pH. This acidification of the endosome lumen is mediated primarily by the vacuolar proton (H^+^) ATPase (V-ATPase), which pumps cytosolic H^+^ into endosomes in association with the hydrolysis of ATP (Ohkuma et al., 1982).

Mitochondria are essential organelles in eukaryotes, serving as a site for ATP synthesis, lipid metabolism, and Ca^2+^ storage (Mesmin, 2016; Rizzuto et al., 2012; Wang and Youle, 2009). In addition to these basic cell homeostatic functions, mitochondria play a key role in apoptosis (Green and Reed, 1998). Apoptotic stimuli, including DNA damage–induced cellular stress, result in translocation of members of the Bcl-2 family of apoptosis regulatory proteins to mitochondria and the consequent release of cytochrome c from these organelles into the cytosol (Kluck et al., 1997). The pore proteins that mediate this release of cytochrome c are voltage-dependent anion channels (VDACs) (Cheng et al., 2003; Shimizu et al., 1999).

The three mammalian members of the VDAC family of proteins (VDAC1 to VDAC3) share ~70% amino acid sequence identity (Amodeo et al., 2014). The two yeast members of this family, YVDAC1 and YVDAC2, are encoded by the *POR1* and *POR2* genes, respectively, and share 50% sequence identity (Blachly-Dyson et al., 1997). All VDAC proteins localize to the outer mitochondrial membrane and possess intrinsic pore-forming activity. Crystal structure analysis revealed that zebrafish VDAC2 forms a pore structure composed of 19 β-strands (Schredelseker et al., 2014). The metabolites that permeate through VDACs include ATP, ADP, NADH, succinate, phosphate, and creatine phosphate (Pavlov et al., 2005; Zalman et al., 1980). The open probability of the VDAC pore is determined by the mitochondrial membrane potential (Vander Heiden et al., 2000), which influences insertion of the NH2-terminal region of the protein into the pore (Schredelseker et al., 2014). The pore diameter of VDAC channels thus changes from 2.5 nm in the open state to 1.8 nm in the closed state in response to a change in membrane potential (Colombini et al., 1987; Mannella, 1987). Although ATP and other metabolites are small enough to pass through the VDAC pore even in the closed state, they are not able to do so (Tan and Colombini, 2007), suggesting that the permeability to ions and metabolites is determined by the electrostatic nature of the inner wall of the pore in addition to the pore diameter (Colombini, 2016).

It has remained unclear whether all VDAC proteins share a common function or whether individual family members have distinct roles. The reduced growth rate of Δ*por1* mutant yeast was rescued by forced expression of mouse VDAC1, VDAC2, or VDAC3 (Dihanich et al., 1987; Sampson et al., 1997), indicating that all mammalian VDACs share the same function. On the other hand, the phenotypes of knockout mice have suggested that VDAC2 may have a specific function whose loss cannot be compensated for by VDAC1 or VDAC3. Mice deficient in VDAC1 or VDAC3 are thus viable but manifest mitochondrial dysfunction (Anflous et al., 2001) and infertility (Sampson et al., 2001), respectively, whereas VDAC2 knockout mice die in utero (Cheng et al., 2003). Mice lacking VDAC2 specifically in the heart are born alive but die shortly thereafter as a result of progressive fibrosis and cardiomyopathy (Raghavan et al., 2012). The specific function of VDAC2 that underlies this difference in phenotypes has remained unknown, however.

We have previously shown that endosomal localization of a complex formed by the small GTPase Ras and PI3K promotes clathrin-independent endocytosis and endosome maturation (Fujioka et al., 2011). In addition, a 28–amino acid sequence at the NH_2_-terminal region of the Ras binding domain (RBD) of PI3K was found to be required for such endosomal localization and was thus designated RAPEL (<u>R</u>as-<u>P</u>I3K <u>e</u>ndosomal localization) (Fujioka et al., 2019). However, the molecular mechanism by which the Ras-PI3K complex is recruited to endosomes in a RAPEL-dependent manner has remained to be investigated. We have now identified the mitochondrial protein VDAC2 as a RAPEL binding protein. We further found that VDAC2, but not VDAC1 or VDAC3, contributes to regulation not only of the endosomal localization of the Ras-PI3K complex but also of subsequent endosome maturation in response to epidermal growth factor (EGF). Moreover, VDAC2 was shown to mediate the tethering of endosomes to mitochondria and thereby to promote endosome maturation. Our study has thus uncovered a specific role for VDAC2 in the regulation of mitochondrion-endosome association and endosome maturation.

## RESULTS

### VDAC2 Interacts with RAPEL of PI3K and Regulates Endosomal Localization of the Ras-PI3K Complex

To reveal the molecular mechanism by which the Ras-PI3K complex is translocated from the plasma membrane to endosomes in a RAPEL-dependent manner (Fujioka et al., 2019), we screened for RAPEL binding proteins with the use of a yeast two-hybrid assay based on the RBD of PI3K (PI3K-RBD) as the bait (**Figure S1A**). We identified six candidate proteins (**Table S1**), among which we focused on the mitochondrial pore protein VDAC2 because we were able to confirm its interaction with RAPEL in mammalian cells by immunoprecipitation analysis (**Figure 1A**).

**Figure 1.**
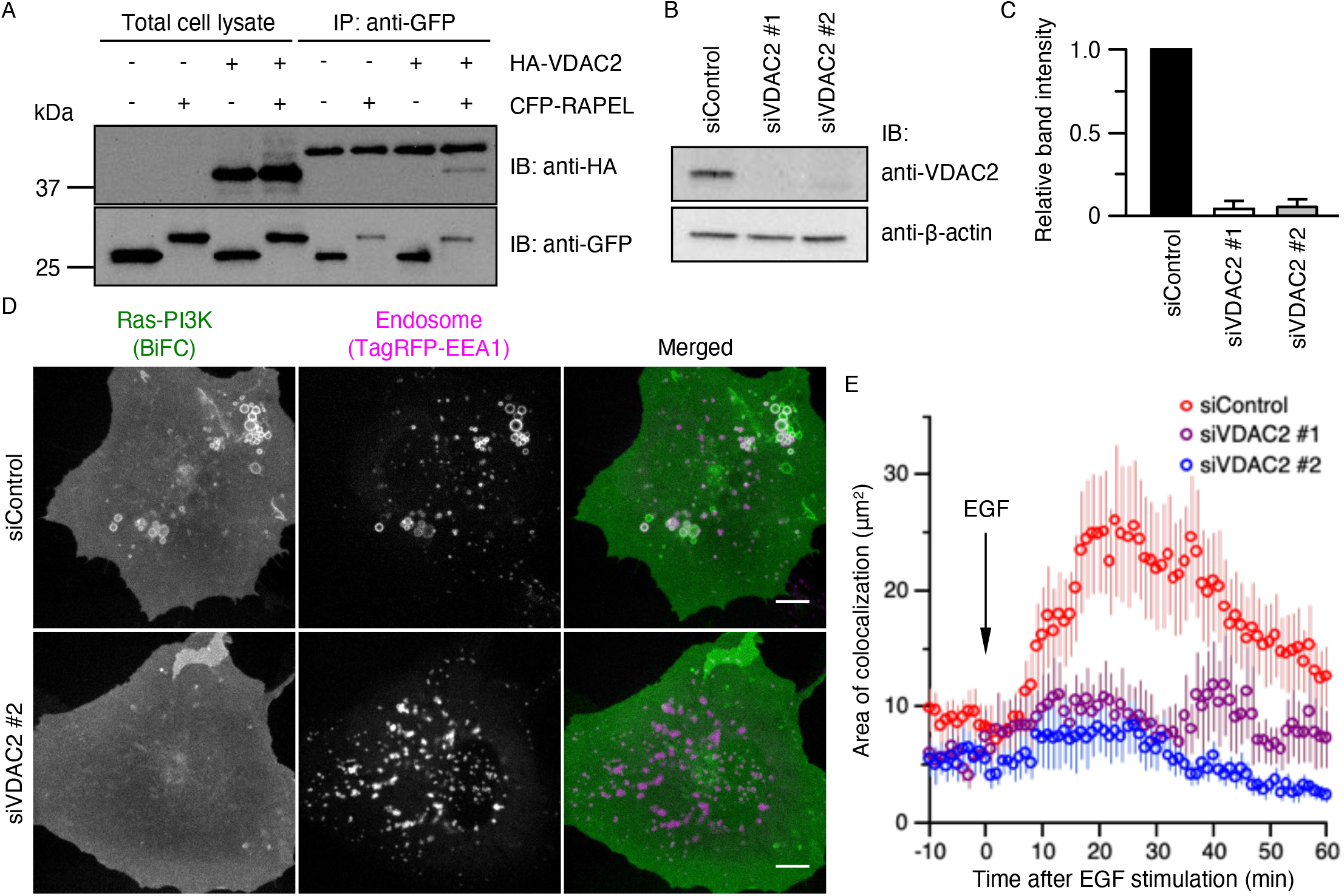
VDAC2 Is a RAPEL Binding Protein Required for Endosomal Localization of the Ras-PI3K Complex. (A) HEK293T cells expressing hemagglutinin epitope (HA)–tagged VDAC2 or cyan fluorescent protein (CFP)–tagged RAPEL, (or CFP alone) were subjected to immunoprecipitation (IP) with antibodies to green fluorescent protein (GFP), and the resulting precipitates as well as a portion of the total cell lysates were subjected to immunoblot analysis (IB) with antibodies to HA and to GFP. (B) Immunoblot analysis of VDAC2 and β-actin (loading control) in A431 cells transfected for 72 h with control (siControl) or VDAC2 (siVDAC2 #1 or #2) siRNAs. (C) Band intensity of VDAC2 normalized by that of β-actin for immunoblots as in (B). Data are means + SEM from three independent experiments. (D) A431 cells were transfected with siControl or siVDAC2 #2 for 48 h and then further transfected with expression vectors for EEA1 (TagRFP-EEA1), H-Ras (VN-H-Ras-WT), and the RBD of PI3K (PI3K-RBD-VC) for 24 h. The cells were then deprived of serum for 4 h and subjected to time-lapse microscopic observation with exposure to EGF (100 ng/ml) at time 0. Representative BiFC (Ras-PI3K complex) and TagRFP (EEA1) images at 30 min after the onset of EGF stimulation are shown. Bars, 10 µm. (E) Time course of colocalization area for the Ras-PI3K complex and EEA1 in images as in (D). Data are means ± SEM for a total of at least 14 cells from three independent experiments. *p* < 0.0001 (MANOVA with Bonferron’s correction). See also Figure S1, Table S1, and Movies S1 and S2.

To examine whether VDAC2 contributes to the EGF-induced endosomal localization of the Ras-PI3K complex, we prepared two independent small interfering RNAs (siRNAs) specific for VDAC2 mRNA (siVDAC2 #1 and siVDAC2 #2). Immunoblot analysis revealed that introduction of each of these siRNAs into A431 cells resulted in an ~90% reduction in the abundance of VDAC2 (**Figures 1B and 1C**). Consistent with previous findings (Tsutsumi et al., 2009), exposure of control cells to EGF resulted in colocalization of the Ras-PI3K complex and the early endosome marker EEA1, which were visualized by bimolecular fluorescence complementation (BiFC) analysis and with the use of TagRFP-labeled EEA1, respectively (**Figures 1D and 1E**; **Movie S1**). In contrast, the EGF-induced translocation of the Ras-PI3K complex from the plasma membrane to endosomes was markedly attenuated in cells depleted of VDAC2 (**Figures 1D and 1E**; **Movie S2**). We also confirmed the interaction of full-length PI3K and VDAC2 by immunoprecipitation analysis in HEK293T cells (**Figure S1B**). In this instance, the interaction between the two proteins was not affected by EGF stimulation, suggesting that it might be constitutive.

Given that VDACs are major components of the outer mitochondrial membrane, we next examined whether knockdown of VDAC2 might affect mitochondrial morphology or the mitochondrial ATP concentration. Despite the fact that they both attenuated endosomal localization of the Ras-PI3K complex (**Figures 1D and 1E**), siVDAC2 #1 induced mitochondrial fragmentation and an increase in the mitochondrial ATP concentration, whereas cells transfected with siVDAC2 #2 did not manifest any pronounced changes in mitochondrial morphology or ATP concentration (**Figures S1C–S1E**). Given that overexpression of VDAC2 also induced mitochondrial fragmentation (**Figure S1F**), we concluded that siVDAC2 #1 might have off-target activity and we therefore used only siVDAC2 #2 in subsequent experiments. Collectively, these results indicated that VDAC2 is a RAPEL binding protein and contributes directly to regulation of the endosomal localization of the Ras-PI3K complex.

### VDAC2 Is a Positive Regulator of Clathrin-independent Endocytosis

To evaluate whether VDAC2 might regulate endocytosis, we incubated VDAC2-depleted cells with fluorescent dye–labeled 10-kDa dextran, which is incorporated into cells via the clathrin-independent endocytosis pathway (Araki et al., 2003). Knockdown of VDAC2 in HEK293T, HeLa, or A431 cells suppressed dextran uptake by ~20% compared with that in corresponding control cells (**Figure 2A**). Conversely, overexpression of VDAC2 promoted dextran uptake in these cell lines (**Figure 2B**), indicating that VDAC2 positively regulates clathrin-independent endocytosis. Knockdown or overexpression of VDAC2 did not affect uptake of transferrin (**Figures 2C and 2D**), which is internalized into cells via clathrin-dependent endocytosis (Harding et al., 1983), suggesting that VDAC2 is dispensable for the regulation of clathrin-dependent endocytosis.

**Figure 2.**
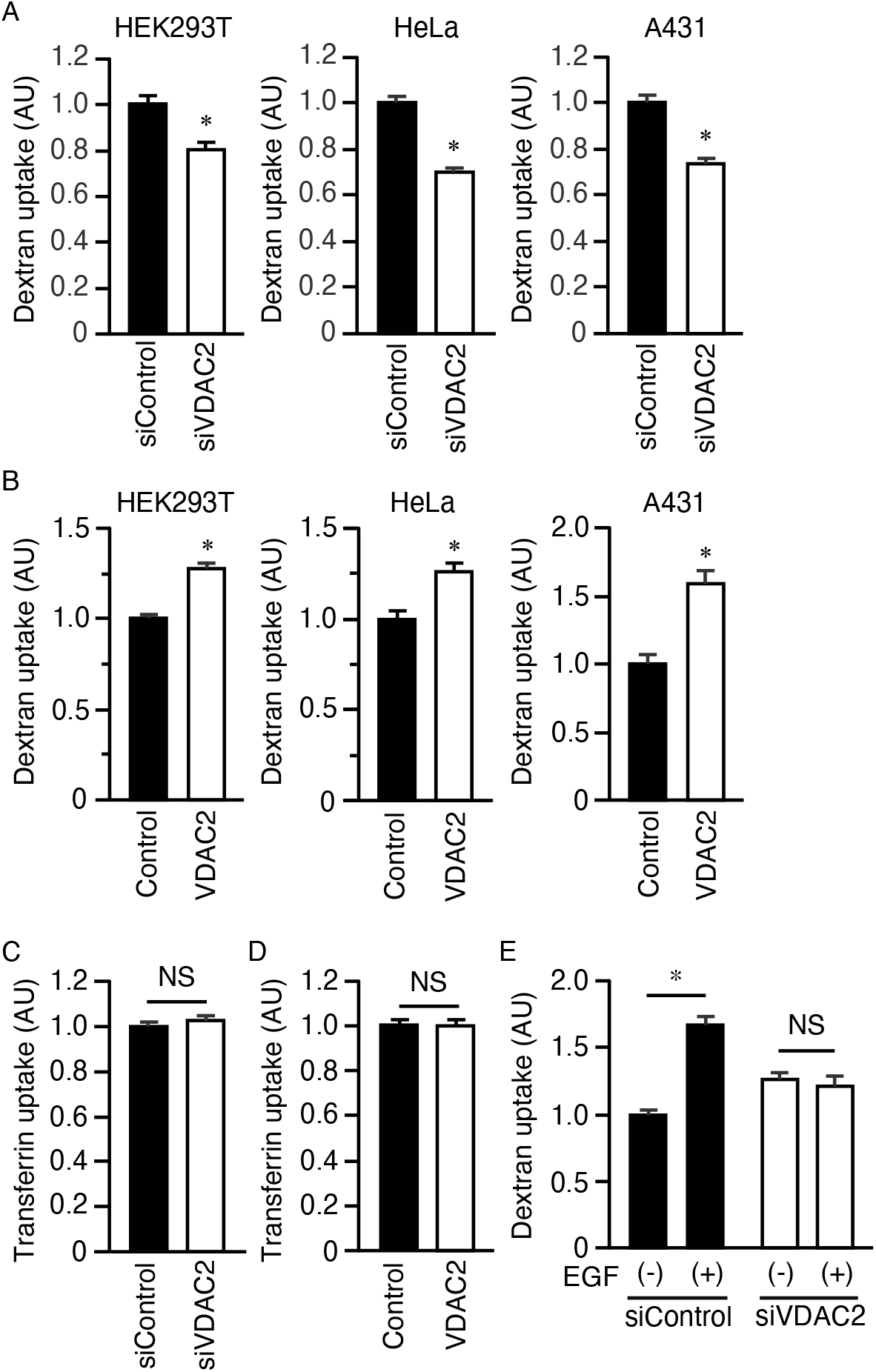
VDAC2 Contributes to Up-regulation of Clathrin-Independent Endocytosis by EGF. (A) HEK293T, HeLa, or A431 cells were transfected with control or VDAC2 siRNAs for 72 h and then incubated with Alexa Fluor 546–labeled dextran for 30 min before determination of total fluorescence intensity (AU, arbitrary units) within the cells. Data are means + SEM for a total of at least 280 (HEK293T), 254 (HeLa), or 201 (A431) cells from three independent experiments. *p < 0.0005 versus siControl-transfected cells (Student’s t test). (B) Dextran uptake in HEK293T, HeLa, or A431 cells expressing the CFP variant SECFP either alone (Control) or together with HA-tagged VDAC2 was quantitated as in (A). Data are means + SEM for a total of at least 302 (HEK293T), 112 (HeLa), or 148 (A431) cells from three independent experiments. \**p* < 0.0005 versus cells transfected with the control vecttor (Student’s t test). (C) HEK293T cells transfected with control or VDAC2 siRNAs for 72 h were incubated with Alexa Fluor 546–labeled transferrin for 30 min before measurement of total fluorescence intensity within the cells. Data are means + SEM for a total of at least 325 cells from three independent experiments. NS, not significant (p = 0.9309, Student’s t test). (D) Transferrin uptake in HEK293T cells expressing SECFP either alone (Control) or together with HA-VDAC2 was quantitated as in (C). Data are means + SEM for a total of at least 277 cells from three independent experiments. NS, p = 0.9868 (Student’s t test). (E) A431 cells were transfected with control or VDAC2 siRNAs for 72 h, deprived of serum for 4 h, and incubated with Alexa Fluor 546–labeled dextran in the absence or presence of EGF (100 ng/ml) for 30 min before measurement of total fluorescence intensity within the cells. Data are means + SEM for a total of at least 220 cells from three independent experiments. \**p* < 0.001 (one-way ANOVA followed by Tukey’s HSD test).

We next investigated the role of VDAC2 in EGF-induced up-regulation of endocytosis. Whereas dextran uptake was significantly increased by EGF stimulation in siControl-transfected A431 cells, consistent with previous findings (Liberali et al., 2008), this effect was abolished in VDAC2-depleted cells (**Figure 2E**). These results thus implicated VDAC2 in the up-regulation of clathrin-independent endocytosis by EGF.

### VDAC2 Mediates EGF-Induced Mitochondrion-Endosome Association

We identified VDAC2 as a RAPEL binding protein and showed that it contributes to the regulation of endocytosis mediated by Ras-PI3K signaling. To explore the mechanism by which the mitochondrial protein VDAC2 participates in the regulation of endocytosis, we first examined the intracellular localization of VDAC2 and the Ras-PI3K complex in detail. We found that a fraction of endosomes positive for the Ras-PI3K complex was localized in close proximity to VDAC2-positive mitochondria in EGF-stimulated A431 cells (**Figures 3A and 3B**). Three-dimensional (3D) reconstruction from confocal images also revealed the apparent association of mitochondria with endosomes positive for the Ras-PI3K complex (**Figure 3C**). Quantitation of this association between mitochondria and endosomes showed it to be increased by EGF in control cells but not in VDAC2-depleted cells (**Figure 3D**). Together, these results suggested that endosomes positive for the Ras-PI3K complex associate with mitochondria through VDAC2 in an EGF-dependent manner.

**Figure 3.**
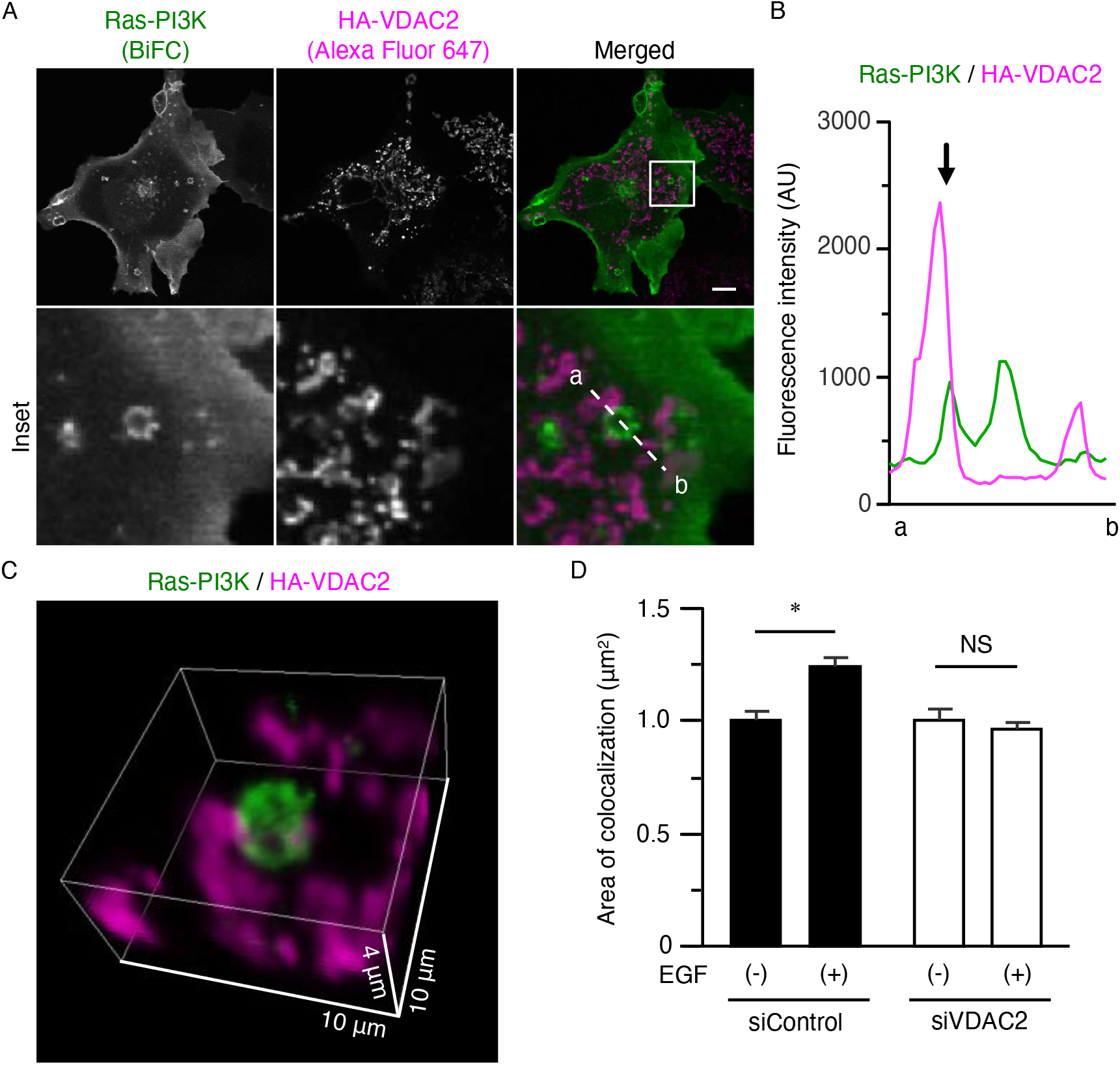
Mitochondrion-Endosome Association Is Promoted by EGF in a VDAC2-Dependent Manner. (A) A431 cells expressing HA-tagged VDAC2 as well as H-Ras (VN-H-Ras-WT) and the RBD of PI3K (PI3K-RBD-VC) were deprived of serum for 4 h, stimulated with EGF (100 ng/ml) for 15 min, fixed, and subjected to immunofluorescence staining with antibodies to HA and Alexa Fluor 647—conjugated secondary antibodies. The localization of the Ras-PI3K complex was also examined by BiFC analysis. The boxed region in the upper merged image is shown at higher magnification in the lower panels. Bar, 10 µpm. (B) Fluorescence intensities of the Ras-PI3K complex (green) and VDAC2 (magenta) along the line indicated by a and b in the inset of the merged image in (A). The arrow indicates contact between a VDAC2-positive mitochondrion and an endosome positive for the Ras-PI3K complex. (C) Representative 3D image reconstructed from confocal images of the cell in (A). Ras-PI3K complex–positive endosomes and VDAC2-positive mitochondria are shown in green and magenta, respectively. (D) A431 cells were transfected with control or VDAC2 siRNAs for 48 h and then further transfected with an expression vector encoding mitochondrially targeted enhanced green fluorescent protein (mito-EGFP) for 24 h. They were then deprived of serum for 4 h, incubated in the absence or presence of EGF (100 ng/ml) for 15 min, fixed, and subjected to immunofluorescence staining with antibodies to Rab5 and Alexa Fluor 647–conjugated secondary antibodies. The area of colocalization of mito-EGFP with Rab5 was quantified. Data are means + SEM for a total of at least 78 determinations from three independent experiments. \**p* < 0.001 (one-way ANOVA followed by Tukey’s HSD test).

### VDAC2 Promotes Endosome Acidification by Mediating Mitochondrion-Endosome Association

Multicolor live-cell imaging revealed that the Ras-PI3K complex is localized in early endosomes (visualized by TagRFP-EEA1) in EGF-stimulated A431 cells (Tsutsumi et al., 2009), and that such endosomes appeared to make contact with mitochondria over a period of several minutes (**Figure 4A** and **Movie S3**). The fluorescence signal for EEA1 gradually decreased during this period (**Figure 4A** and **Movie S3**), suggesting the possibility that VDAC2 might promote endosome maturation, in addition to its regulation of clathrin-independent endocytosis, through interorganelle association. To examine this possibility more directly, we monitored endosomal pH in VDAC2-overexpressing cells with the use of dextran labeled with the pH-sensitive fluorescent dye AcidiFluor ORANGE. Given that overexpression of VDAC2 promotes dextran uptake (**Figure 2B**), dextran labeled with (pH-insensitive) Alexa Fluor 488 was also included as a reference. The ratio of AcidiFluor ORANGE to Alexa Fluor 488 fluorescence intensities was determined to reflect normalized pH in endosomes, and it was found to be increased in VDAC2-overexpressing cells (**Figures 4B and 4C**). We also used another pH-sensitive dye, the ECGreen-Endocytosis Detection reagent (ECGreen), which stains the plasma membrane and endosomes, in order to track individual endosomes for direct comparison of endosomal pH between before and after endosome association with mitochondria in the presence of EGF. The fluorescence intensity of ECGreen was significantly increased after mitochondrion-endosome association compared with before (**Figures 4D and 4E**; **Movie S4**). Furthermore, this increase was abrogated by VDAC2 knockdown (**Figures 4D and 4E**; **Movie S5**). The fluorescence intensity of the pH-insensitive dye PlasMem Bright Red, which stains the plasma membrane, was not affected by mitochondrion-endosome association (**Figure S2A**). Collectively, these results suggested that VDAC2 promotes endosome acidification by mediating mitochondrion-endosome association.

**Figure 4.**
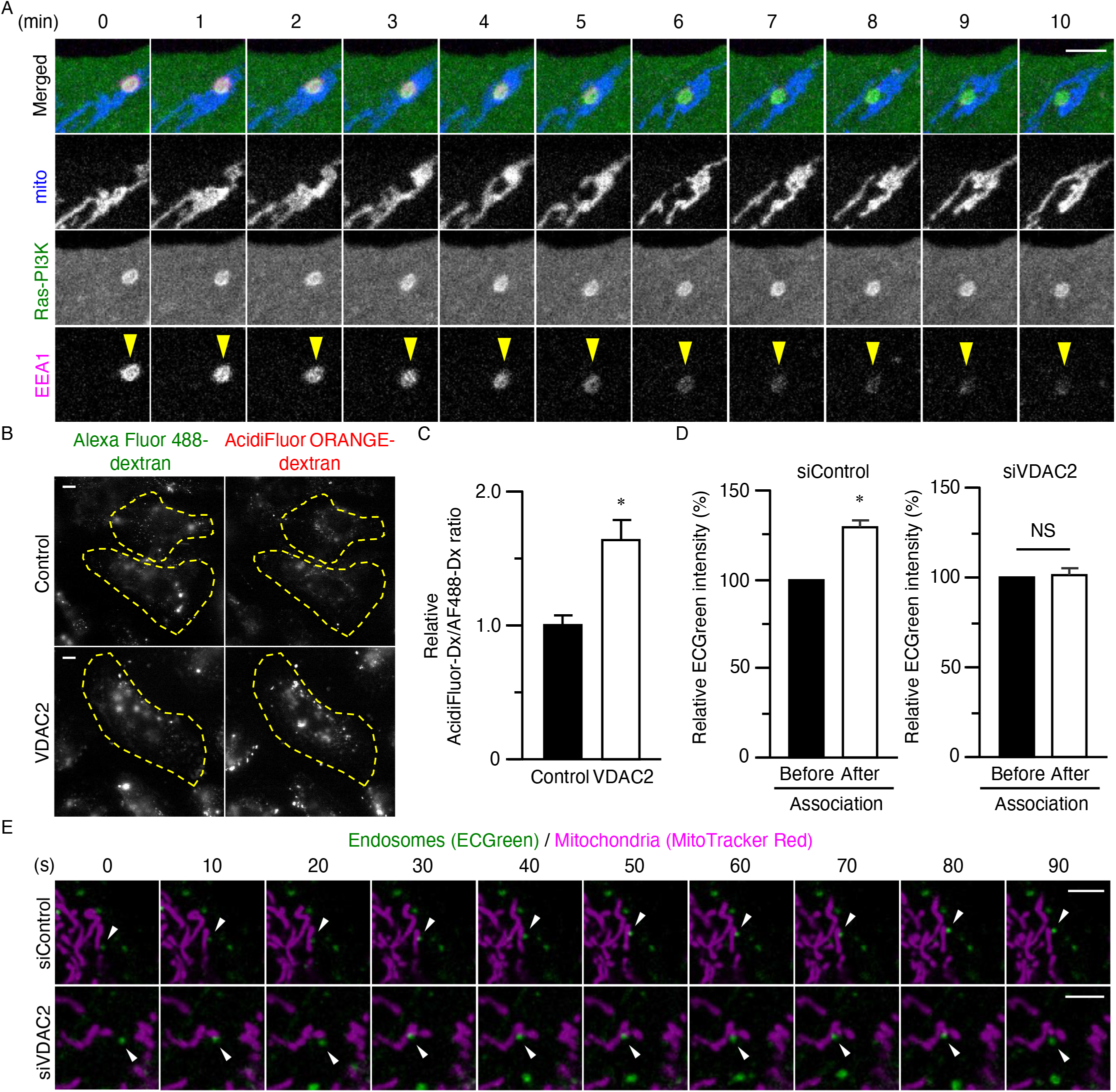
Endosome Acidification Is Promoted by Mitochondrion-Endosome Association. (A) A431 cells transfected with expression vectors for mitochondrially targeted SECFP (mito-SECFP), VN-H-Ras-WT, PI3K-RBD-VC, and TagRFP-EEA1 were deprived of serum for 4 h and then stimulated with EGF (100 ng/ml). Representative time-lapse images over 10 min are shown. The arrowhead indicates an endosome positive for Ras-PI3K complex whose fluorescence intensity of EEA1 was dicreased during the association with mitocondria. Bar, 5 µm. (B) HeLa cells expressing SECFP either alone (Control) or together with HA-tagged VDAC2 were incubated with Alexa Fluor 488–labeled dextran and AcidiFluor ORANGE–labeled dextran for 15 min before imaging by fluorescence microscopy. Yellow dashed lines indicate individual cells expressing SECFP. Bars, 10 µm. (C) Ratio of the fluorescence intensity of AcidiFluor ORANGE–labeled dextran (Dx) to that of Alexa Fluor 488–labeled dextran for cells as in (B). Data ares means + SEM for a total of at least 81 cells from three independent experiments. \**p* < 0.0005 versus control vector–transfected cells (Student’s t test). (D) A431 cells transfected with control or VDAC2 siRNAs for 72 h were deprived of serum for 4 h, incubated first with MitoTracker Red CMXRos for 20 min and then with ECGreen for 10 min, and then stimulated with EGF (100 ng/ml). The average fluorescence intensity of individual ECGreen-positive vesicles was determined before and after their EGF-induced association with mitochondria. Data are means + SEM for a total of 30 cells from three independent experiments. \**p* < 0.0001 versus value for before association with mitochondria (Wilcoxon signed-rank test). (E) Representative time-lapse images of siControl- or siVDAC2-transfected cells over 90 s in the presence of EGF as in (D). Arrowheads indicate ECGreen-positive endosomes. Bars, 5 µm. See also Figure S2 and Movies S3 to S5.

Given that VDAC family members have been found to functionally compensate for the loss of other isoforms (Sampson et al., 1997), we tested whether VDAC1 or VDAC3 might also contribute to the regulation of endosome acidification. Although all three mammalian VDAC isoforms interacted with RAPEL (**Figure S2B**), the acidification of endosomes after their association with mitochondria was still apparent in A431 cells after specific knockdown of VDAC1 or VDAC3 (**Figures S2C and S2D**). These results supported the notion that, among the mammalian VDAC isoforms, VDAC2 specifically contributes to endosome acidification by mediating mitochondrion-endosome association.

### VDAC2 Tethers Mitochondria to Endosomes and Contributes Functionally to the Regulation of Endocytosis

Finally, we evaluated whether the mitochondrion-endosome association per se promotes endosome acidification. To this end, we established an optogenetics system to induce such association by blue light illumination. Cryptochrome 2 (CRY2) is a blue light photoreceptor of *Arabidopsis thaliana* that undergoes a conformational change in response to illumination at a wavelength of 488 nm. This change in conformation promotes the binding of CRY2 to cryptochrome-interacting basic helix-loop-helix (CIB1). We therefore constructed expression vectors for modified forms of CRY2 (CRY2-iRFP-FYVE or CRY2-MiCy-FYVE) and CIB1 (TOM20-mCherry-CIB1 or TOM20-iRFP-CIB1) that localize to endosomes and mitochondria, respectively (**Figure 5A**). Exposure of A431 cells transfected with these vectors to light at 488 nm indeed markedly increased the colocalization of CRY2-iRFP-FYVE and TOM20-mCherry-CIB1 (**Figures 5B and 5C**). This illumination also significantly increased endosome acidification in such cells as assessed with the use of AcidiFluor ORANGE–labeled dextran (**Figure 5D**). This light-induced endosome acidification was not observed in VDAC2-depleted cells (**Figure 5E**), indicative of a role for VDAC2 in endosome acidification other than its mediation of mitochondrion-endosome association. Blue illumination did not induce a significant change in the fluorescence intensity of dextran labeled with the pH-insensitive dye Alexa Fluor 546 in such a system (**Figure S3A**).

**Figure 5.**
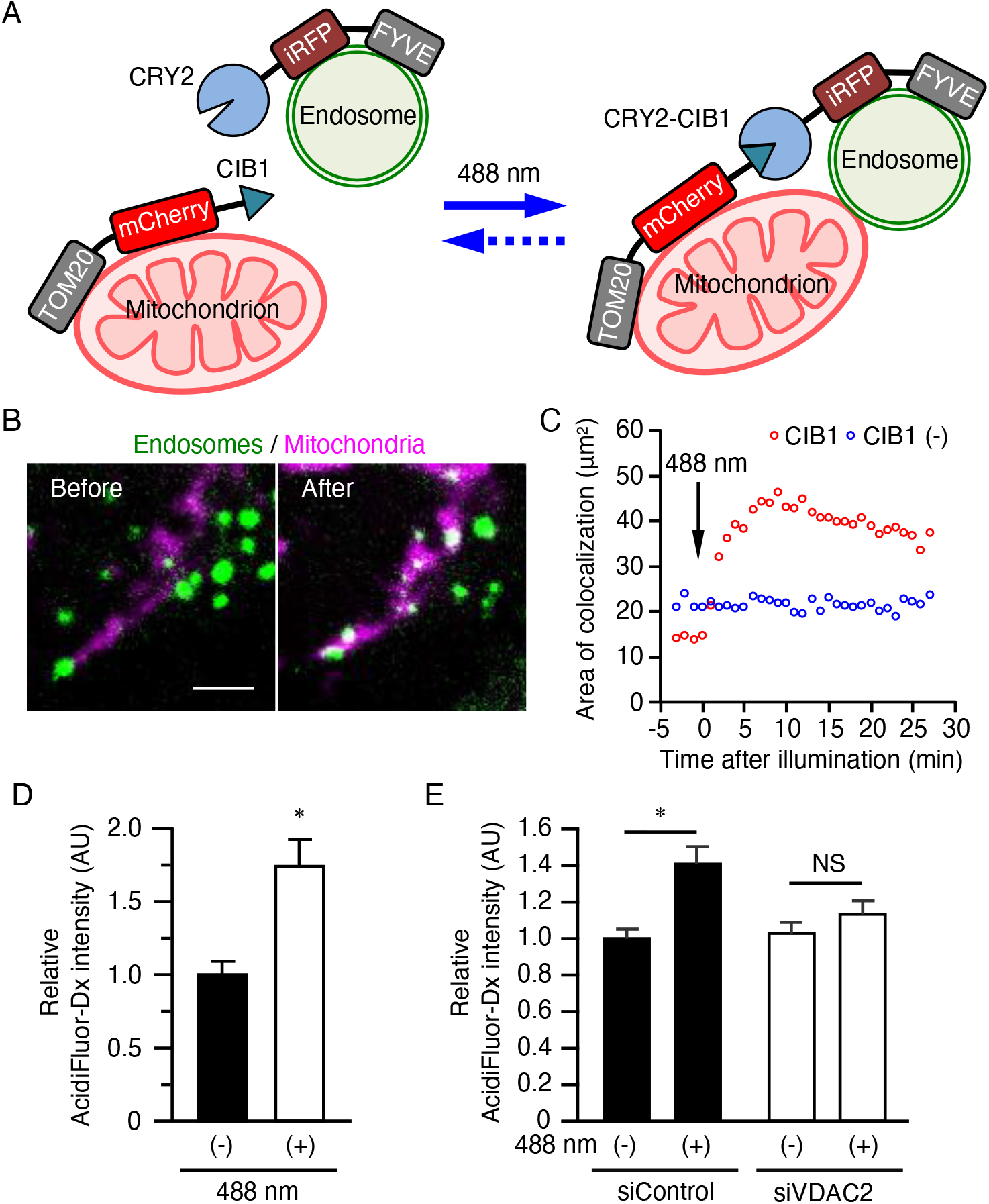
VDAC2 Is Required for the Promotion of Endosome Acidification Dependent of Mitochondrion-Endosome Association. (A) Schematic representation of optogenetic manipulation of mitochondrion-endosome association. Endosome-associated CRY2-iRFP-FYVE binds to mitochondrially localized TOM20-mCherry-CIB1 in a manner dependent on illumination at a wavelength of 488 nm. (B) A431 cells transfected with expression vectors for CRY2-iRFP-FYVE and TOM20-mCherry-CIB1 were subjected to time-lapse microscopy and exposed to 488-nm light for 5 s during imaging. Representative images of the distribution of mitochondria (magenta) and endosomes (green) before and after the illumination are shown. Bar, 5 µm. (C) Time course of the area of colocalization of mitochondria and endosomes determined as in (B). Cells expressing CRY2-iRFP-FYVE and TOM20-mCherry [CIB1 (–)] were also examined as a negative control. Data are means ± SEM for a total of at least 10 cells from three independent experiments. p < 0.0001 (MANOVA). (D) A431 cells expressing CRY2-MiCy-FYVE and TOM20-iRFP-CIB1 were incubated with AcidiFluor ORANGE–labeled dextran (AcidiFluor-Dx) for 15 min before illumination at 488 nm for 5 s and imaging by fluorescence microscopy. Total fluorescence intensity of AcidiFluor-Dx within the cells was quantitated. Data ares means + SEM for a total of at least 39 cells from three independent experiments. \**p* < 0.0001 versus nonilluminated cells (Student’s t test). (E) A431 cells were transfected with siControl or siVDAC2 for 48 h and then further transfected with expression vectors for CRY2-MiCy-FYVE and TOM20-iRFP-CIB1 for 24 h. They were then incubated with AcidiFluor-Dx for 15 min before illumination at 488 nm for 5 s and imaging by fluorescence microscopy. Total fluorescence intensity of AcidiFluor-Dx within the cells was quantitated. Data are means + SEM for a total of at least 39 cells from three independent experiments. \**p* < 0.01 (one-way ANOVA followed by Tukey’s HSD test). See also Figure S3.

We further investigated the function of VDAC2 that is required for endosome acidification. VDAC proteins were originally identified as anion channels, but the pore of these proteins is now known to allow the passage of cations and metabolites (Pavlov et al., 2005; Zalman et al., 1980). Moreover, a mutant of yeast VDAC1 with reduced permeability to ions and small molecules has been established (Blachly-Dyson et al., 1990; Ellenrieder et al., 2019). We therefore constructed a permeability-deficient mutant of human VDAC2 (K31E, K72E) on the basis of sequence similarity with yeast VDAC1. The acidity of endosomes as reflected by the fluorescence intensity of LysoTracker was increased by expression of an siRNA-resistant form of wild-type VDAC2 in A431 cells depleted of endogenous VDAC2 (**Figure S3B**). Similar forced expression of the permeability-deficient mutant of VDAC2 also induced a small increase in endosomal acidity, but this effect was significantly less pronounced than was that induced by wild-type VDAC2 (**Figure S3B**). These results thus implicated the pore function of VDAC2 in the promotion of endosome acidification.

## DISCUSSION

We have here identified VDAC2, a mitochondrial outer membrane protein, as a RAPEL binding protein and revealed that it contributes to endosomal localization of the Ras-PI3K complex as well as to subsequent clathrin-independent endocytosis and endosome maturation. In this context, VDAC2 functions not only as a tethering factor for endosomes and mitochondria but also as a pore protein for the permeation of ions or metablites. Our findings thus uncover a previously unappreciated role for VDAC2 in the regulation of endosome maturation dependent on interorganelle association between endosomes and mitochondria.

Endosomes are known to associate with other organelles including the endoplasmic reticulum (ER) (Rowland et al., 2014) and mitochondria (Das et al., 2016; Sheftel et al., 2007). The first indication of contact between ER and endocytic organelles was provided by a study of yeast (Pan et al., 2000), with evidence of ER-endosome association subsequently being obtained in mammalian cells (Eden et al., 2010; Rocha et al., 2009). Although the composition of membrane contact sites between ER and endosomes remains to be fully characterized, vesicle-associated membrane protein (VAMP)–associated proteins (VAPs) are thought to function as tethering factors (Alpy et al., 2013; Dong et al., 2016; Rocha et al., 2009). Proposed functions of endosome-ER association include determination of endosome fate between the degradative and recycling pathways (Rowland et al., 2014), regulation of growth factor receptor signaling (Eden et al., 2010), cholesterol transport from endocytosed low-density lipoproteins to ER (Du et al., 2011), and bidirectional Ca^2+^ mobilization (Van Der Kant and Neefjes, 2014). Previously described functions of endosome-mitochondrion association have been limited to unidirectional transport from endosomes to mitochondria. For example, the interaction of endosomes containing transferrin with mitochondria results in transfer of transferrin-bound iron from the former to the latter organelles (Das et al., 2016). Such iron transfer might play an important role in mitochondrial function given that iron is a constituent of the active center of enzymes that catalyze redox reactions (Hentze et al., 2004) as well as of heme and iron-sulfur clusters (Braymer and Lill, 2017). We have now shown that mitochondria associate with endosomes in a manner dependent on the interaction of VDAC2 with PI3K, and that they actively regulate endosome maturation. The association of endosomes with mitochondria, like that with ER, therefore also has bidirectional functional consequences.

How does the mitochondrial protein VDAC2 regulate endosome maturation? In addition to H^+^, other ions including Na^+^, K^+^, Cl^-^, and Ca^2+^ have been implicated in endosome maturation (Scott and Gruenberg, 2011). Given that VDACs have been shown to transport Ca^2+^, K^+^, Cl^-^, and ATP (Báthori et al., 2006; Peng et al., 1992; Rostovtseva and Colombini, 1996; Zambrowicz and Colombini, 1993) between mitochondria and the cytosol, factors that are required for endosome maturation might be transferred from mitochondria to endosomes via VDAC2. Endosome maturation is accompanied by hydrolysis of cytosolic ATP by V-ATPase and consequent transport of H^+^ across the endosomal membrane to render the lumen acidic (Ohkuma et al., 1982). Mitochondria might therefore supply ATP via VDAC2 at contact sites with endosomes in order to maintain the local ATP concentration for V-ATPase. Given that the mitochondrial ATP concentration (~2 mM) is lower than that in the cytosol (~8 mM) (Imamura et al., 2009), however, a mechanism to overcome this unfavorable concentration gradient might be required for the movement of ATP from mitochondria to the cytosol. Mammalian VDAC1, through interaction with the inositol 1,4,5-trisphosphate (IP3) receptor, participates in Ca^2+^ transport from ER to mitochondria at contact sites between these organelles (Szabadkai et al., 2006). Moreover, the accumulation of Ca^2+^ in endosomes is required for the heterotypic fusion of late endosomes and lysosomes (Wong et al., 2017). Ca^2+^ transport mediated by VDAC2 at mitochondrion-endosome contact sites, similar to that at ER-mitochondrion contact sites, might therefore contribute to endosome maturation. In addition to Ca^2+^, Cl^-^ may play a role in endosome maturation. Acidification of the endosomal lumen as a result of the proton pump activity of V-ATPase might be expected to be accompanied by anion influx. For example, in osteoclasts, the transport of Cl^-^ into the endosomal lumen by the H^+^/Cl^-^ exchange transporter CLC7 facilitates efficient proton pumping by V-ATPase (Kornak et al., 2001). Although the Cl^-^ concentration in mitochondria has not been precisely determined, Cl^-^ might be released from mitochondria into the cytosol through VDAC2 at contact sites between mitochondria and endosomes. Such released Cl^-^ may then be transported into endosomes by an unknown transporter in the endosomal membrane in order to maintain electroneutrality in the endosomal lumen. Consistent with this notion, expression of the permeability-deficient mutant of VDAC2 was only partially able to support endosome acidification compared with the wild-type protein (**Figure S3B**). However, the partial ability of the VDAC2 mutant in this regard suggests that an unknown function of VDAC2 not dependent on its permeability also contributes to endosome maturation.

The mammalian VDAC family comprises three isoforms that share the function of a mediator of ion and metabolite flux between mitochondria and the cytosol (Raghavan et al., 2012). However, mice deficient in VDAC2, but not those deficient in VDAC1 or VDAC3, die during embryogenesis (Cheng et al., 2003), suggesting that VDAC2 has a unique role during embryonic development whose loss cannot be compensated for by other VDAC members. We have now found that VDAC2, but not VDAC1 or VDAC3, contributes to the regulation of endocytosis by mediating mitochondrion-endosome association in an EGF-dependent manner. Given that endocytosis contributes to the fine-tuning of signal transduction through internalization of cell surface receptors, it might be expected that signaling pathways subject to strict spatiotemporal control during embryogenesis become aberrantly activated in mice lacking specific endocytosis regulators (Seto et al., 2002). For instance, mice deficient in Vam2, a component of the homotypic fusion and vacuole protein sorting (HOPS) complex, manifest embryonic mortality, and endosome maturation from late endosomes to lysosomes was found to be suppressed in embryonic fibroblasts established from these animals (Aoyama et al., 2012). This study further implicated an endocytic anomaly that gives rise to sustained activation of bone morphogenetic protein signaling in embryonic mortality (Aoyama et al., 2012). Moreover, EGF has been found to play a role in the regulation of oocyte maturation and embryonic development (Lonergan et al., 1996). VDAC2 might therefore be essential for embryogenesis as a result of its contribution to the regulation of endocytosis by EGF.

In conclusion, we have shown that direct association between mitochondria and endosomes promotes endosome maturation. Such association is mediated by VDAC2, which also contributes to endosome maturation in a manner independent of its role as a tethering factor. Future studies, including visualization of the local dynamics of ions and ATP at mitochondrion-endosome contact sites, are warranted to unveil the molecular mechanism underlying this function of VDAC2.

## ACKNOWLEDGMENTS

We thank A. Miyawaki for providing Venus and SECFP cDNAs, L. Stephen{s?} for p110γ cDNA, M. Matsuda for rabbit antiserum to GFP, T. Tsuboi for pEGFP-C1-Rab5, H. Noji for the mitoATeam1.03 vector, T. Miyazaki for pGBKT7 and pGADT7, and K. Aoki for pCXpuro-CRY2-cRaf, as well as A. Kikuchi for technical assistance. This work was supported in part by Grants-in-Aid from the Ministry of Education, Culture, Sports, Science, and Technology of Japan (18H0485009, 20H0489500, and 20H20H05872) and from Japan Society for the Promotion of Science (15K152305, 16H0622709, 17H0401609, 19KK0198, 19H04823, and 20K1611500).

## AUTHOR CONTRIBUTIONS

Conceptualization: Y.O. Methodology: A.O.S., S.K., Y.F., and Y.O. Formal analysis: A.O.S., S.K., and Y.F. Investigation: A.O.S. and Y.F. Resources: M.F., Y.F., S.K., and H.S. Writing–original draft: A.O.S., Y.F., and Y.O. Writing–review and editing: S.K., A.Y., Y.F., H.S., A.N., and Y.O. Visualization: A.O.S. and Y.F. Supervision: Y.F., H.S., A.N., M.A., and Y.O. Project administration: Y.O. Funding acquisition: A.O.S., Y.F., and Y.O.

## STAR Methods

### Reagents and Antibodies

Recombinant human EGF was obtained from PeproTech (Rocky Hill, NJ, USA). Alexa Fluor 488– or Alexa Fluor 546–labeled dextran (10 kDa), Alexa Fluor 546–labeled transferrin, Alexa Fluor 647–labeled antibodies to rat or mouse immunoglobulin G (IgG), MitoTracker Red CMXRos, MitoTracker Green FM, and LysoTracker Red DND-99 were obtained from Thermo Fisher Scientific (Carlsbad, CA, USA). AcidiFluor ORANGE–labeled dextran was from Goryo Chemical (Sapporo, Japan). ECGreen-Endocytosis Detection and PlasMem Bright Red reagents were obtained from Dojindo Laboratories (Mashiki Japan). Protein G– or protein A–conjugated Sepharose beads were from GE Healthcare (Little Chalfont, UK). Horseradish peroxidase (HRP)–conjugated goat antibodies to rat, mouse, or rabbit IgG were obtained from Jackson ImmunoResearch Laboratories (West Grove, PA, USA), and HRP-conjugated donkey antibodies to goat IgG were from Promega (Madison, WI, USA). Antibodies to GFP and V5, to HA (3F10), to β-actin, to VDAC2, and to Rab5 were obtained from Medical & Biological Laboratories (Nagoya, Japan), Thermo Fisher Scientific, Roche (Indianapolis, IN, USA), Santa Cruz Biotechnology (Santa Cruz, CA, USA), Atlas Antibodies (Stockholm, Sweden), and BD Biosciences (San Joe, CA, USA), respectively. Rabbit antiserum to GFP was kindly provided by M. Matsuda (Kyoto University).

### Cell Culture and Transfection

HEK293T (CRL011268), A431 (CRL-1555), and HeLa (CCL-2) cells were obtained from American Type Culture Collection (Manassas, VA, USA) and were cultured under a humidified atmosphere containing 5% CO2 at 37°C in Dulbecco’s modified Eagle’s medium (DMEM) (Sigma-Aldrich, Tokyo, Japan) supplemented with 10% fetal bovine serum (Thermo Fisher Scientific). Expression vectors were introduced into HEK293T and A431 cells by transfection for 24 h with the use of Polyethylenimine Max (Polysciences, Warrington, PA, USA), and into HeLa cells by transfection for 24 h with the use of FuGENE HD (Promega).

### Plasmids

The expression vectors pCAGGS-VN-H-Ras-WT, pCXN2-FLAG-PI3K(p110γ)-RBD-VC, pFX-SECFP, pFX-ECFP-RAPEL, pFX-mito-SECFP, and pFX-mito-EGFP were described previously (Fujioka et al., 2018, 2019; Kashiwagi et al., 2019; Tsutsumi et al., 2009). The vector pCMV-TagRFP-EEA1 was obtained from Addgene (Cambridge, MA, USA). Expression vectors for EGFP-Rab5 and mitoATeam1.03 (Imamura et al., 2009) were kindly provided by T. Tsuboi (The University of Tokyo) and H. Noji (University of Tokyo), respectively. The vectors pGBKT7 and pGADT7 were kindly provided by T. Miyazaki (Hokkaido University). The vector pCXN2-FLAG-H-Ras-WT (Ohba et al., 2000) was digested with NotI, rendered blunt-ended with Klenow enzyme, and further digested with EcoRI after phenol-chloroform extraction to obtain the coding sequence of human H-Ras-WT. The resulting DNA fragment was subcloned into the EcoRI and SmaI sites of pGADT7 to obtain pGADT7-H-Ras-WT. The coding sequence for the RBD of PI3K (human p110γ) was amplified from pCXN2-FLAG-PI3K-RBD-VC by the polymerase chain reaction (PCR) with the primer pair PI3KRBD_F and PI3KRBD_R. The resulting PCR product was subcloned into the EcoRI and SalI sites of pGBKT7 to obtain pGBKT7-PI3KRBD. Human cDNAs for VDAC1, VDAC2, and VDAC3 were generated from a human 293T cDNA library by PCR with the primer pairs VDAC1_F and VDAC1_R, VDAC2_F and VDAC2_R, and VDAC3_F and VDAC3_R, respectively. These sequences were then subcloned into the XhoI and NotI sites of pCXN2-3×HA or pCXN2-FLAG-H-Ras-WT-IRES-EGFP to obtain pCXN2-3×HA-VDAC and pCXN2-FLAG-VDAC-IRES-EGFP vectors, respectively. The coding sequence for an siRNA-resistant form of VDAC2 was obtained by PCR-based mutagenesis with the use of a QuikChange Site-Directed Mutagenesis Kit (Agilent Technologies, Santa Clara, CA, USA) and with the primer pair siVDAC2#2_F and siVDAC2#2_R, and a cDNA encoding the K31E-K72E mutant of siRNA-resistant VDAC2 was also obtained by PCR-based mutagenesis with the primer pairs K31E_F and K31E_R as well as K72E_F and K72E_R. These PCR products were subcloned into the XhoI and NotI sites of pCXN2-FLAG-H-Ras-WT-IRES-EGFP. The coding sequence of the V5 tag was generated by annealing of the oligonucleotide pair V5tag_F and V5tag_R, and was then subcloned into the EcoRI and BglII sites of pFXII-EGFP-Rab5 to obtain pFXII-V5-Rab5. The coding sequence of PI3K was cleaved out of pCXN2-FLAG-PI3K-RBD-VC by digestion with XhoI and NotI and was subcloned into the EcoRI and BglII sites of pFXII-V5-Rab5 to obtain pFXII-V5-PI3K. The coding sequence of *A. thaliana* CIB1 was amplified from pCIB1(deltaNLS)-pmGFP (Addgene) by PCR with the primer pair CIB1_F and CIB1_R and was then subcloned into the BamHI and BglII sites of pFX-TOM20-mCherry or pFX-TOM20-iRFP (Kashiwagi et al., 2019) to obtain pFX-TOM20-mCherry-CIB1 and pFX-TOM20-iRFP-CIB1, respectively. The cDNA for *A. thaliana* CRY2 was released from pCX4puro-CRY2-cRaf (kindly provided by K. Aoki, National Institutes for Basic Biology, Okazaki, Japan) by digestion with EcoRI and XhoI and was subcloned into the corresponding sites of pFX-FYVE to obtain pFX-CRY2-FYVE. The cDNAs for MidoriishiCyan (MiCy) and infrared fluorescent protein (iRFP) were subcloned into the NotI and BglII sites of pFX-CRY2-FYVE to obtain pFX-CRY2-MiCy-FYVE and pFX-CRY2-iRFP-FYVE, respectively. All PCR products were confirmed by sequencing analysis, and all primer sequences for vector construction are provided in **Table S2**.

### Yeast Two-Hybrid Assay

A yeast two-hybrid assay was performed with the use of the Matchmaker Gold Yeast Two-Hybrid System (Clontech-Takara, Kusatsu, Japan). In brief, the Y2H Gold strain was transformed with pGBKT7-PI3KRBD, which encodes the DNA binding domain of Gal4 fused to PI3K-RBD (Gal4-BD–PI3K-RBD). The Y187 strain was transformed with pGADT7-H-Ras-WT, which encodes the activation domain of Gal4 fused to wild-type human H-Ras (H-Ras-WT). Control experiments were performed with the following combinations before screening: pGBKT7-p53 and pGADT7-T (positive control), pGBKT7-PI3KRBD and pGADT7-H-Ras-WT (positive control), and pGBKT7-Lam and pGADT7-T (negative control). After testing the bait for autoactivation and toxicity, Y2H Gold cells expressing Gal4-BD–PI3K-RBD and Y187 cells transformed with a human cDNA library [Mate & Plate Library–Universal Human (Normalized), Clontech] were cocultured for 22 h at 30°C in YPDA medium, after which mating of the cells was confirmed by phase-contrast microscopy. The mated cells were collected by centrifugation, resuspended in 0.5× YPDA medium, diluted with 0.9% NaCl (to 1:10, 1:100, 1:1000, and 1:10,000), and then streaked on SD/–Trp, SD/–Leu, SD/–Trp/–Leu (double dropouts, DDO), and DDO/X-α-gal (X)/Aureobasidin A (A) plates. After incubation for 5 days at 30°C, colonies that had formed on the DDO/X/A plates were streaked on an SD/–Ade/–His/–Leu/–Trp/X/A (quadruple, QDO/X/A) plate and incubated at 30°C. A total of 57 colonies was collected and cultured in YPDA medium, and the cells were subsequently isolated by centrifugation at 600 x *g* for 30 s. The cell pellets were suspended in SZB buffer [0.5 M EDTA (pH 8.0), 1 M sodium citrate, 2 M sorbitol, 0.5% Zymolyase, 0.7% 2-mercaptoethanol], incubated for 40 min at 37°C, and, after the addition of potassium acetate to a final concentration of 1.5 M, incubated for an additional 5 min on ice. After centrifugation of the cell lysates at 20,000 x *g* for 15 min at 4°C, portions (300 µl) of the supernatants were mixed with 3 M sodium acetate (200 µl) and 2-propanol (1 ml), incubated for 10 min at −20°C, and centrifuged at 20,000 x *g* for 5 min at 4°C. Yeast plasmids were recovered by the ethanol precipitation method and used to transform *Escherichia coli* (JM109 strain). The plasmids were subsequently purified from the bacterial cells with the use of a FastGene Plasmid Mini Kit (Nippon Genetics, Tokyo, Japan) and subjected to DNA sequencing analysis with the use of a T7-promoter sequencing primer.

### Immunoblot Analysis

Unless otherwise specified, cells were lysed in NP-40 lysis buffer [10 mM Tris-HCl (pH 7.4), 150 mM NaCl, 5 mM EDTA, 0.5% Nonidet P-40, 10% glycerol, 1 mM NaF, 1 mM Na_3_VO_4_, 1 mM phenylmethylsulfonyl fluoride] by incubation for 30 min on ice. The lysates were centrifuged at 20,000 x *g* for 10 min at 4°C, and the resulting supernatants were subjected to SDS-polyacrylamide gel electrophoresis. The separated proteins were transferred to a polyvinylidene difluoride membrane (Millipore, Darmstadt, Germany) and subjected to immunoblot analysis. Nonspecific sites on the membrane were blocked by incubation for 1 h with 5% dried skim milk before exposure overnight at 4°C to primary antibodies (anti-GFP and anti-HA, 1:1000 dilution; anti-VDAC2, 1:500; anti-V5, 1:5000). The membrane was washed three times with Tris-buffered saline containing 0.05% Tween 20 (TBS-T) and then incubated for 1 h at room temperature with HRP-conjugated secondary antibodies (1:5000). After washing of the membrane an additional three times with TBS-T, immune complexes were detected with the use of ECL Western Blotting Detection Reagent (GE Healthcare) and a MIIS imaging system (Givetechs, Sakura, Japan).

### Immunoprecipitation

Cell lysates prepared as for immunoblot analysis were incubated with primary antibodies for 1 h at 4°C with gentle rotation, after which protein A– or protein G–Sepharose beads were added and the incubation was continued for an additional 1 h. Immune complexes bound to the beads were isolated by centrifugation, washed three times with NP-40 lysis buffer, and eluted with SDS sample buffer. After heating at 95°C for 5 min, the samples were subjected to immunoblot analysis.

### Fluorescence Microscopy

Cell images were acquired with IX83 or IX81 (Olympus, Tokyo, Japan) or Eclipse *Ti2* (Nikon, Tokyo, Japan) microscopes. Both IX83 and IX81 microscopes were equipped with a BioPoint MAC 6000 filter and shutter control unit (Ludl Electronic Products, Hawthorne, NY, USA), an automated XY-stage (Chuo Precision Industrial, Tokyo, Japan), a UPlanSApo 60×/1.35 oil objective lens, and a SOLA Light Engine (Lumencor, Beaventon, OR, USA). The Eclipse T*i*2 microscope was equipped with a PlanApo 10×/0.45 objective lens and an X-Cite Turbo multiwavelength LED illumination system (Excelitas Technologies, Waltham, MA, USA). The IX83, IX81, and Eclipse T*i*2 microscopes were also equipped with a Rolera EM-C^2^ electron-multiplying cooled charge-coupled device (CCD) camera (QImaging, Surrey, British Columbia, Canada), a Zyla 4.2 scientific complementary metal-oxide semiconductor (sCMOS) camera (Andor Technology, Belfast, UK), and a Zyla 5.5 sCMOS camera (Andor Technology), respectively. Representative excitation and emission filters adopted for this study included FF-02-438/24 and FF01-483/32 (Semrock, Rochester, NY, USA) or CFP HQ (Nikon) for CFP; BP470-490 and BP510-550 (Olympus) for GFP and Alexa Fluor 488; FF01-500/24-25 and FF01-542/27 (Semrock) for YFP (BiFC signal); FF02-438/24 and FF01-542/27 for FRET; and BP520-550 and BA580IF (Olympus) or Cy3 HQ (Nikon) for TagRFP, Alexa Fluor 546, and Alexa Fluor 594. MetaMorph software (Molecular Devices, San Jose, CA, USA) was used for the control of microscopes and peripheral equipment. For live-cell imaging, cells were incubated at 37°C with a Chamlide incubator system (Live Cell Instrument, Seoul, Korea). Confocal images were acquired with an FV10i confocal microscope (Olympus) and either an sDISK (Andor Technology) or X-Light V2 (CrestOptics, Rome, Italy) spinning-disk confocal unit.

### Quantification of Endosomal Localization of Ras-PI3K

Analysis of the endosomal localization of the Ras-PI3K complex was performed as described previously (Fujioka et al., 2019). In brief, confocal images of cells expressing Venus NH2–terminus-Ras (VN-H-Ras-WT), PI3K-RBD–Venus COOH-terminus (PI3K-RBD-VC), and TagRFP-EEA1 were acquired, and background signals were subtracted. Vesicle structures in BiFC images (Ras-PI3K complex) and TagRFP images (EEA1) were respectively extracted with the use of the ‘Top Hat’ module of MetaMorph software. Colocalization area for the Ras-PI3K complex and EEA1 was quantified with the use of the ‘measure colocalization’ function of the software.

### RNA Interference

The siRNAs specific for human VDAC1 mRNA (siVDAC1, s14768), VDAC2 mRNA (siVDAC2 #1, s14773; siVDAC2 #2, s14772), or VDAC3 mRNA (siVDAC3, s230730) were obtained from Ambion (Austin, TX, USA). Cells were transfected 72 h with siRNAs with the use of the Lipofectamine RNAiMAX reagent (Thermo Fisher Scientific). AllStars Negative Control siRNA (Qiagen, Valencia, CA, USA) was used as a control siRNA (siControl).

### RNA Isolation and RT-qPCR Analysis

Total RNA was purified from cells with the use of an RNeasy Mini Kit (Qiagen), and portions (2.5 µg) were subjected to reverse transcription (RT) with the use of a SuperScript VILO cDNA Synthesis Kit (Thermo Fisher Scientific). The resulting cDNA was subjected to quantitative PCR (qPCR) analysis with the primers listed in Table S3.

### Quantification of Mitochondrial Fragmentation and ATP Concentration

For analysis of mitochondrial fragmentation, A431 cells plated on collagen-coated glass-bottom dishes (35-mm diameter; Matsunami Glass, Kishiwada, Japan) were incubated with MitoTracker Red CMXRos (12.5 nM) for 10 min at 37°C and then washed three times with phosphate-buffered saline (PBS) before image acquisition by sDISK spinning-disk confocal microscopy. The extent of mitochondrial fragmentation was quantitated with ImageJ software according to previously described protocols (Giedt et al., 2016; Rehman et al., 2012). In brief, background-subtracted images were subjected to a median filter and processed with the ImageJ plugin ‘Auto local threshold’ in order to generate mitochondrial mask images. Mitochondrial particles were counted with the ImageJ plugin ‘Particle counting.’ The mitochondrial fragmentation count [in arbitrary units (AU)] was calculated from the mask images as the number of mitochondrial particles x 10,000/total area of mitochondria (pixels). For analysis of mitochondrial ATP concentration, A431 cells were transfected with the mitoATeam1.03 vector for 24 h before imaging with the IX83 wide-field fluorescence microscope. The Förster resonance energy transfer (FRET)/CFP emission ratio was calculated from background-subtracted images as described previously (Mizutani et al., 2010).

### Endocytosis Assay

For evaluation of clathrin-independent or clathrin-dependent endocytosis, cells plated on collagen-coated glass-bottom dishes (35-mm diameter) were incubated with Alexa Fluor dye–conjugated dextran (10-kDa) or transferrin at 500 µg/ml for 30 min at 37°C and then washed three times with PBS before image acquisition with the IX83 wide-field fluorescence microscope. After background subtraction, total fluorescence intensity within cells was quantified with MetaMorph software.

### Immunofluorescence Analysis

Cells were fixed with 3% paraformaldehyde for 15 min, permeabilized with 0.1% Triton X-100 in PBS for 4 min, and then exposed to 1% bovine serum albumin for 30 min at room temperature before incubation first overnight at 4°C with primary antibodies (anti-HA or anti-Rab5, 1:1000 dilution) and then for 1 h at room temperature in the dark with Alexa Fluor 647–conjugated secondary antibodies (1:250). The cells were imaged with the FV10i confocal microscope (Olympus).

### Analysis of Mitochondrion-Endosome Association

A431 cells seeded on collagen-coated glass-bottom dishes (35-mm diameter) were transfected with pFX-mito-EGFP for 24 h, deprived of serum for 4 h, stimulated with EGF (100 ng/ml) for 15 min, fixed, and subjected to immunofluorescence analysis to visualize endogenous Rab5. The cells were imaged with the FV10i confocal microscope. For quantification of mitochondrion-endosome association, the regions of mitochondria and endosomes were extracted from background-subtracted images with the use of the ‘Top Hat’ module of MetaMorph software. The colocalization area of mitochondria and endosomes was then determined with the use of the ‘measure colocalization’ function of the software.

### Assay of Endosome Maturation

Three independent reagents [AcidiFluor ORANGE–conjugated dextran, LysoTracker Red DND-99, and ECGreen-Endocytosis Detection ([ECGreen)] were adopted for assay of endosome maturation. HeLa cells plated on collagen-coated glass-bottom dishes (35-mm diameter) were incubated with Alexa Fluor 488–conjugated dextran (500 µg/ml) and AcidiFluor ORANGE–conjugated dextran (200 µg/ml) for 15 min at 37°C and then washed three times with ice-cold PBS before image acquisition with the IX83 wide-field microscope. After background subtraction, total fluorescence intensity within the cells was quantitated with MetaMorph software. The extent of endosome acidification was expressed as the ratio of the fluorescence intensities of AcidiFluor ORANGE (pH-sensitive dye) and Alexa Fluor 488 (pH-insensitive dye). Alternatively, A431 cells were incubated with LysoTracker Red DND-99 (10 nM) for 30 min at 37°C and washed three times with ice-cold PBS before imaging with the IX83 wide-field microscope. Total fluorescence intensity within cells was quantitated after background subtraction. In another approach, A431 cells were incubated with MitoTracker Red CMXRos (12.5 nM) for 20 min at 37°C, washed with PBS, incubated with ECGreen (1:1000 dilution) for 10 min at 37°C, and washed again with PBS before stimulation with EGF (100 ng/ml) and analysis by time-lapse confocal microscopy. After background subtraction, the fluorescence intensity of ECGreen of the vesicular structures before and after contact with mitochondria was quantitated.

### Optogenetic Induction of Mitochondrion-Endosome Association

A431 cells were seeded on collagen-coated glass-bottom dishes (35-mm diameter), transfected with pFX-CRY2-iRFP-FYVE and pFX-TOM20-mCherry-CIB1 for 24 h, and subjected to time-lapse confocal microscopy. After 5 min, the cells were exposed to blue light (488 nm) for 5 s in order to induce mitochondrion-endosome association. The regions of mitochondria and endosomes were extracted from the background-subtracted images with the use of the ‘Top Hat’ module of MetaMorph software. The colocalization area of mitochondria and endosomes was then quantified with the use of the ‘measure colocalization’ function of the software. For assay of endosome maturation, A431 cells expressing CRY2-MiCy-FYVE and TOM20-iRFP-CIB1 were incubated with AcidiFluor ORANGE–conjugated dextran (200 µg/ml) for 10 min at 37°C, washed with ice-cold PBS, illuminated with 488-nm light for 5 s, and imaged with the IX83 wide-field microscope. Images were subjected to background subtraction, and the fluorescence intensity of AcidiFluor ORANGE was quantified.

## Statistical Analysis

Data are presented as mean + SEM values from at least three independent experiments (unless indicated otherwise) and were compared between two conditions with Student’s t test or the Mann-Whitney U test, or among more than two conditions by one-way analysis of variance (ANOVA) followed by Tukey’s honestly significant difference (HSD) test. The Wilcoxon signed-rank test was applied for comparison between paired samples. Time-series data sets were compared between two conditions by multivariate analysis of variance (MANOVA), or among more than two conditions by MANOVA with Bonferroni’s correction. No statistical methods were used to predetermine sample size. Studies were performed unblinded. A*p* value of < 0.05 was considered statistically significant, and all statistical analysis was performed with JMP Pro software (version 14.0.0).

## SUPPLEMENTAL MOVIE LEGENDS

**Movie S1. Time-Lapse Imaging of the Ras-PI3K Complex in Control siRNA–Transfected Cells Stimulated with EGF**

A431 cells transfected with siControl for 48 h and then with expression vectors for TagRFP-EEA1, VN-H-Ras-WT, and PI3K-RBD-VC for 24 h were subjected to time-lapse microscopic observation with exposure to EGF at time 0 (related to **Figure 1D**).

**Movie S2. Time-Lapse Imaging of the Ras-PI3K Complex in VDAC2 siRNA–Transfected Cells Stimulated with EGF**

A431 cells transfected with siVDAC2 (#2) for 48 h and then with expression vectors for TagRFP-EEA1, VN-H-Ras-WT, and PI3K-RBD-VC for 24 h were subjected to time-lapse microscopic observation with exposure to EGF at time 0 (related to **Figure 1D**).

**Movie S3. Endosome Acidification Promoted by Mitochondrion-Endosome Association**

A431 cells transfected with expression vectors for mito-SECFP, VN-H-Ras-WT, PI3K-RBD-VC, and TagRFP-EEA1 were deprived of serum for 4 h before stimulation with EGF for 15 min and time-lapse microscopic observation (related to **Figure 4A**).

**Movie S4. Time-Lapse Imaging of Endosome Acidification Dependent on Mitochondrion-Endosome Association in Control siRNA–Transfected Cells**

A431 cells were transfected with siControl for 72 h, deprived of serum for 4 h, incubated first with MitoTracker Red CMXRos for 20 min and then with ECGreen for 10 min, stimulated with EGF, and subjected to time-lapse microscopic observation (related to **Figure 4E**).

**Movie S5. Time-Lapse Imaging of Endosome Acidification Dependent on Mitochondrion-Endosome Association in VDAC2 siRNA–Transfected Cells**

A431 cells were transfected with siVDAC2 for 72 h, deprived of serum for 4 h, incubated first with MitoTracker Red CMXRos for 20 min and then with ECGreen for 10 min, stimulated with EGF, and subjected to time-lapse microscopic observation (related to **Figure 4E**).

## SUPPLEMENTAL TABLES

**Table S1.** **RAPEL-Interacting Proteins Identified by a Yeast Two-Hybrid Assay**

| Protein | GenBank<br>accession number | Protein description |
| --- | --- | --- |
| RNF2 | NM_007212.4 | E3 ubiquitin-protein ligase |
| ATP1B1 | NM_001677.4 | Noncatalytic component of Na<sup>+</sup>/K<sup>+</sup>-ATPase |
| ZNF350 | NM_021632.4 | Transcriptional repressor |
| VDAC2 | NM_001184823.1 | Mitochondrial outer membrane pore protein |
| ANAPC13 | NM_015391.4 | Component of anaphase-promoting complex/cyclosome |
| FAM81B | NM_152548.3 | Unknown |

**Table S2.**
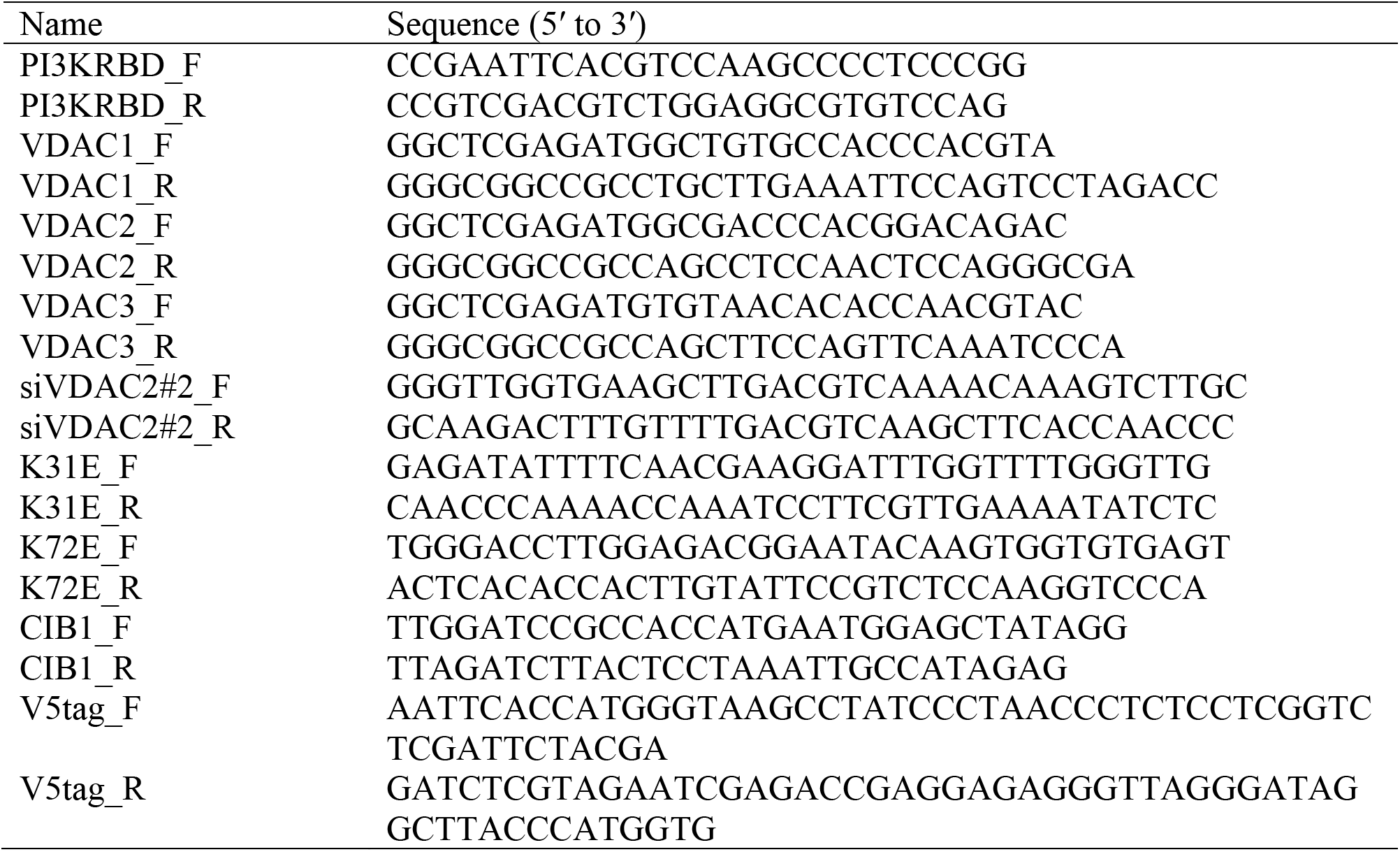
**Primers Used for Vector Construction**

**Table S3.**
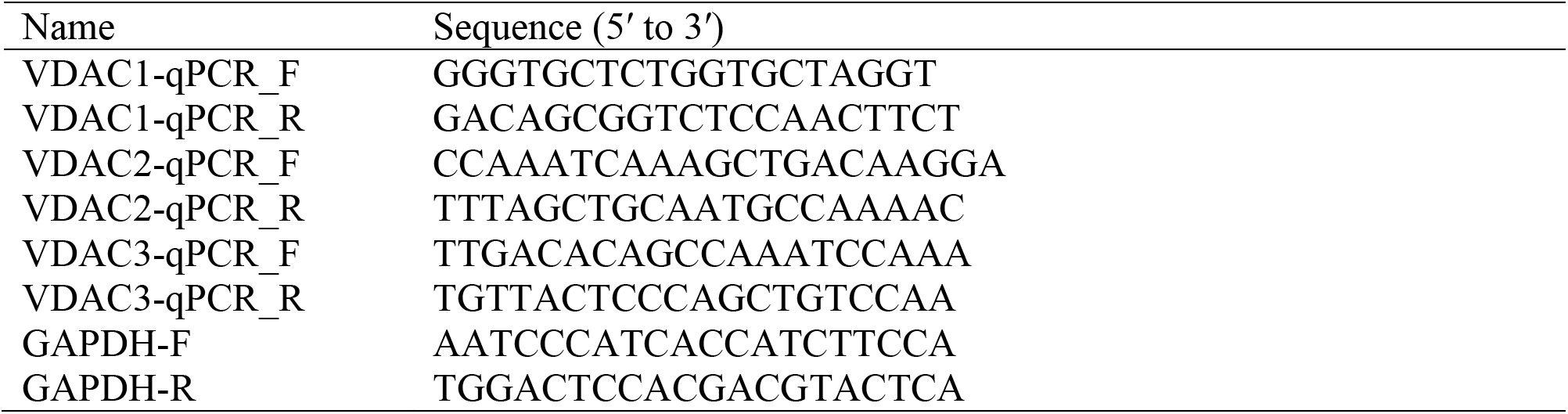
**Primers Used for qPCR Analysis**

## PI3K-VDAC2 interaction tethers endosome and mitochondria to promote endosome maturation

### Graphical Abstract

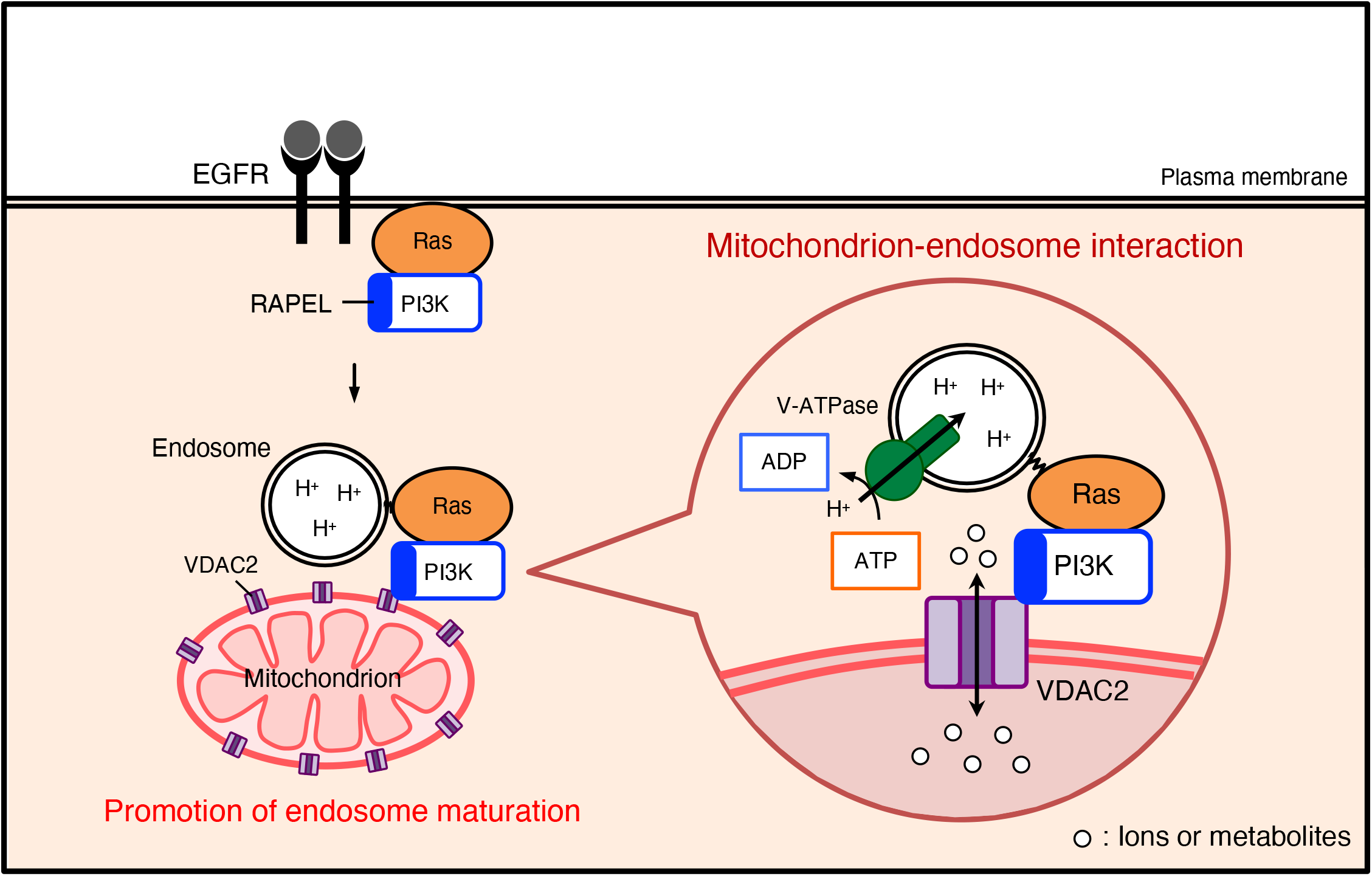

---

**Figure S1.**
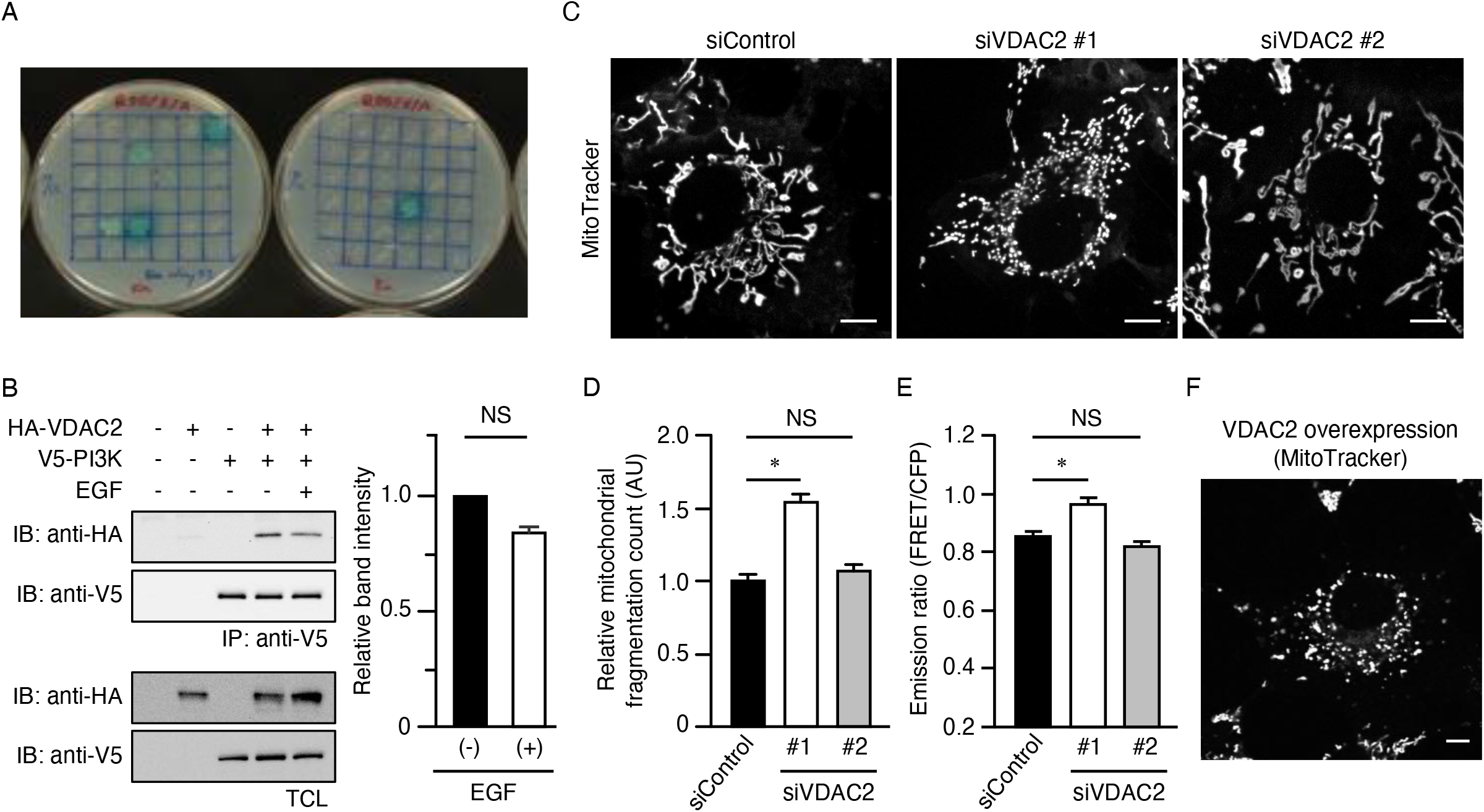
Knockdown of VDAC2, a RAPEL Binding Protein, Does Not Affect Mitochondrial Morphology or ATP Concentration. (A) Yeast two-hybrid assay. Y2H Gold cells transformed with an expression vector encoding the RBD of PI3K fused with the DNA binding domain of Gal4 were mated with Y187 cells transformed with a cDNA library cloned in a vector encoding the activation domain of Gal4 and were then cultured on QDO/X/A plates for 5 days. See Table S1 for a list of PI3K-RBD binding proteins identified. (B) HEK293T cells transfected with expression vectors for HA-tagged VDAC2 or V5-tagged PI3K were deprived of serum for 4 h and then incubated in the absence or presence of EGF (100 ng/ml) for 15 min. The cells were then lysed and subjected to immunoprecipitation (IP) with antibodies to V5, and the resulting precipitates as well as a portion of the total cell lysates (TCL) were subjected to immunoblot (IB) analysis with antibodies to HA and to V5. The band intensity of HA-VDAC2 in immunoprecipitates of cells expressing both exogenous proteins (lane 4 and 5) was determined as the mean + SEM from three independent experiments. NS, not significant (Mann-Whitney U test). (C) A431 cells were transfected with control (siControl) or VDAC2 (siVDAC2 #1 or #2) siRNAs for 72 h, incubated with MitoTracker Red CMXRos for 10 min, and then washed before imaging by confocal microscopy. Bars, 5 µm. (D) Mitochondrial fragmentation count (AU, arbitrary units) for cells as in (C). Data are means + SEM for a total of at least 66 cells from three independent experiments. *p < 0.0001 (one-way ANOVA followed by Tukey’s HSD test). (E) A431 cells were transfected with siControl or siVDAC2 for 48 h and then further transfected with an expression vector for mitoATeam1.03 for 24 h before imaging by fluorescence microscopy. The emission ratio (FRET/CFP) for mitochondria, which reflects the mitochondrial ATP concentration, was determined as mean + SEM values for a total of at least 76 cells from three independent experiments. *p < 0.0005 (one-way ANOVA followed by Tukey’s HSD test). (F) A431 cells transfected with an expression vector for FLAG-tagged VDAC2 and enhanced green fluorescent protein (EGFP) were incubated with MitoTracker Red CMXRos for 10 min, washed, and imaged by confocal microscopy. Bar, 5 µm.

**Figure S2.**
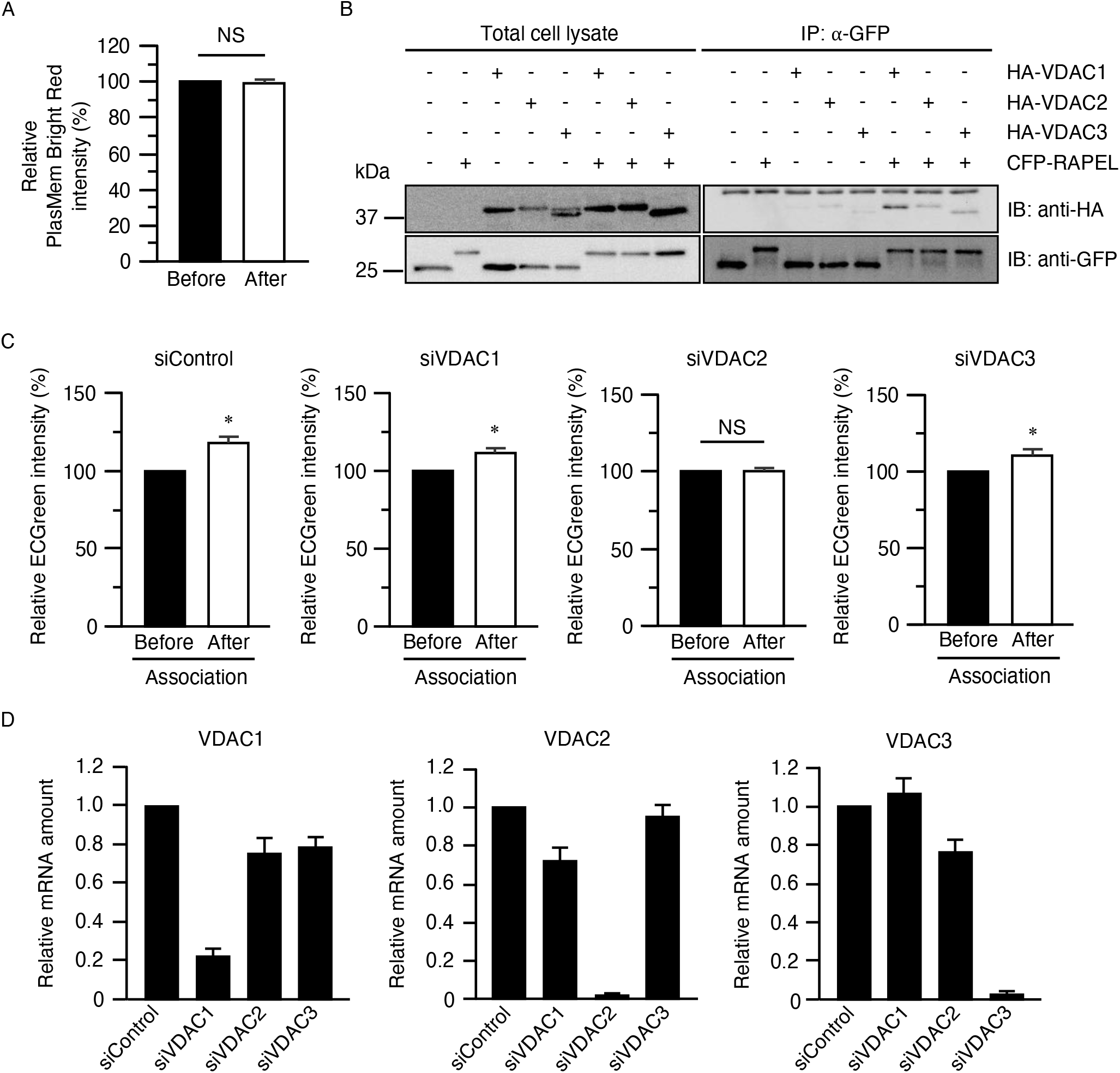
VDAC1 and VDAC3 Are Dispensable for the Promotion of Endosome Acidification by Mitochondrion-Endosome Association. (A) A431 cells were deprived of serum for 4 h, incubated first with MitoTracker Green FM for 20 min and then with PlasMem Bright Red for 5 min, and stimulated with EGF (100 ng/ml) for 20 min. The cells were monitored by time-lapse imaging, and the average fluorescence intensity of individual PlasMem Bright Red–positive vesicles was determined before and after their EGF-induced association with mitochondria. Data are means + SEM for a total of 30 cells from three independent experiments. NS, Wilcoxon signed-rank test. (B) HEK293T cells expressing HA-tagged VDAC1, VDAC2, or VDAC3 together with CFP-tagged RAPEL were subjected to immunoprecipitation (IP) with antibodies to GFP, and the resulting precipitates as a well as a portion of the total cell lysates were subjected to immunoblot analysis (IB) with antibodies to HA and to GFP. (C) A431 cells were transfected with control, VDAC1, VDAC2, or VDAC3 siRNAs for 72 h, deprived of serum for 4 h, incubated first with MitoTracker Red CMXRos for 20 min and then with ECGreen for 10 min, and stimulated with EGF (100 ng/ml). The average fluorescence intensity of individual ECGreen-positive vesicles was measured before and after their EGF-induced association with mitochondria. Data are means + SEM for a total of 30 cells from three independent experiments. *p < 0.05 versus the value before vesicle association with mitochondria (Wilcoxon signed-rank test). (D) Reverse transcription and quantitative polymerase chain reaction analysis of the amount of VDAC1, VDAC2, and VDAC3 mRNAs in A431 cells as in (C). Data were normalized by the amount of glyceraldehyde-3-phosphate dehydrogenase (GAPDH) mRNA and are means + SEM from three independent experiments.

**Figure S3.**
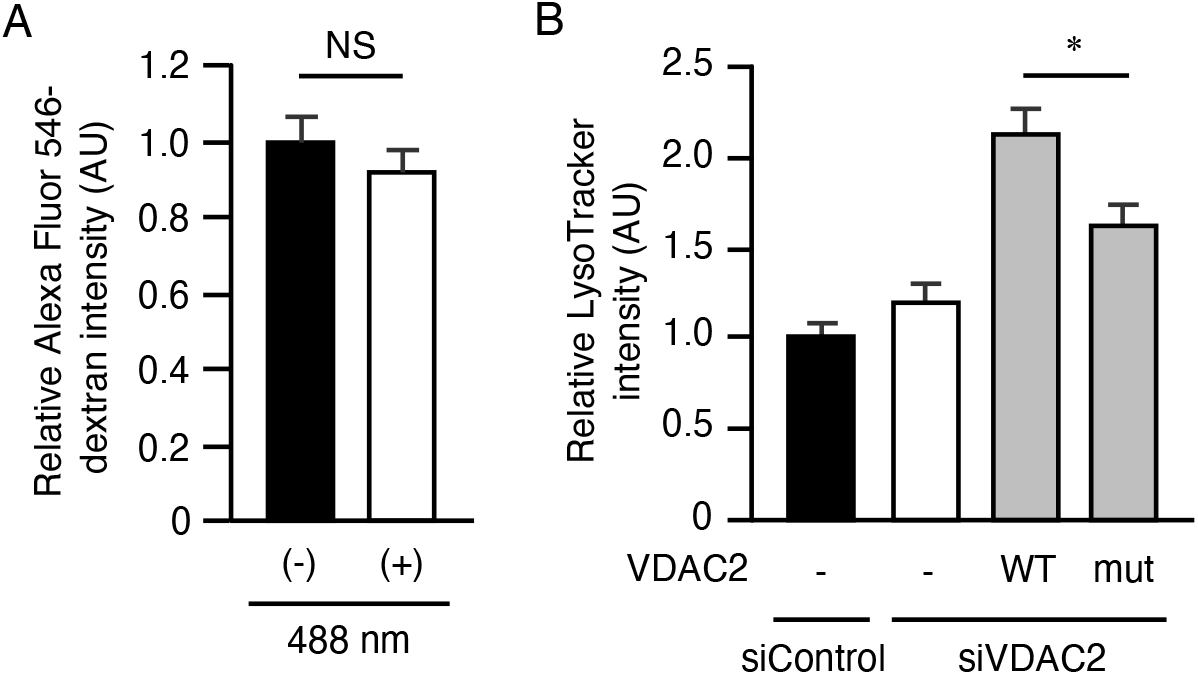
Permeability of VDAC2 Is Partially Required for Endosome Acidification. (A) A431 cells expressing CRY2-MiCy-FYVE and TOM20-iRFP-CIB1 were incubated with Alexa Fluor 546–labeled dextran for 10 min before illumination at 488 nm for 5 s and imaging by fluorescence microscopy. Total fluorescence intensity (AU, arbitrary units) of Alexa Fluor 546–labeled dextran within the cells was quantitated. Data are means + SEM for a total of 64 cells from three independent experiments. NS, Student’s t test. (B) A431 cells were transfected with siControl or siVDAC2 for 48 h and then further transfected with expression vectors for EGFP alone (control) or FLAG-tagged VDAC2-IRES-EGFP [either wild-type (WT) or mutant (K31E, K72E)] for 24 h. The cells were then incubated with 10 nM LysoTracker Red DND-99 for 30 min before fixation and imaging with a fluorescence microscope. Total fluorescence intensity of LysoTracker within the cells was quantitated. Data are means + SEM for a total of at least 105 cells from three independent experiments. *p < 0.0001 (one-way ANOVA followed by Turkey’s HSD test).

